# Insulin Growth Factor 1 affects glutamate receptor activity differently in primary cultures of neocortical versus hippocampal neurons

**DOI:** 10.64898/2026.04.04.716504

**Authors:** Urooj Fatima, Akshay Padala, Steven W. Barger

**Affiliations:** Department of Geriatrics, University of Arkansas for Medical Sciences, Little Rock AR 72205, USA; Department of Neuroscience, University of Arkansas for Medical Sciences, Little Rock AR 72205, USA; Geriatric Research Education & Clinical Center, Central Arkansas Veterans Healthcare System, Little Rock AR 72205, USA

## Abstract

Insulin-like growth factor-1 (IGF-1) plays a critical role in neuronal signaling. Disrupted insulin/IGF-1 signaling is implicated in Alzheimer’s disease, among other conditions, yet its specific influence on glutamate receptor-mediated calcium responses remains unclear. We examined the impacts of IGF-1 on glutamate receptor function in primary rat neurons monitored for intraneuronal calcium following stimulation with glutamate, AMPA, or NMDA/glycine. Pharmacological blockers (CNQX for AMPA receptors, APV for NMDA receptors, and nimodipine for L-type calcium channels) were applied to define receptor-specific contributions. In hippocampal neurons, IGF-1 and insulin altered responses to glutamate in different directions, with IGF-1 tending to evoke and enhanced response. In neocortical neurons, by contrast, IGF-1 consistently reduced glutamate- and AMPA-evoked calcium peaks, suggesting an inhibitory effect on AMPA receptors. To rule out effects on voltage-gated calcium channels downstream of AMPA receptors, we tested effects of IGF-1 on depolarization with potassium chloride; calcium elevation in this case was unaffected by IGF-1. Likewise, IGF-1 did not inhibit responses to NMDA/glycine; and IGF-1 did not affect glutamate responses in the presence of CNQX, a selective AMPA receptor blocker. These findings, combined with the observation that IGF-1 effects persisted in the presence of APV (an NMDA receptor antagonist), indicate that the inhibition of glutamate responses by IGF-1 is mediated by suppression of AMPA receptor activity. IGF-1 may thus contribute to normal neurophysiology, and given the role that glutamate receptors play in excitotoxicity, IGF-1 may confer neuroprotection in the neocortex. Disruption of IGF-1 signaling, as seen in states resembling insulin resistance, may therefore worsen glutamate-driven excitotoxicity and contribute to adverse outcomes.

## INTRODUCTION

Insulin resistance is a common contributing factor to age-related metabolic derangement, compromising glucose uptake into several tissue types and producing type-2 diabetes mellitus (T2DM) in its extreme. While accumulation of glucose in the blood has detrimental consequences, the most severe effects may result from the reduction in this vital energy source in insulin-responsive tissues. Such deficits have been hypothesized to contribute to reductions in cerebral glucose metabolic rates in the development of cognitive impairments and/or dementia **(Baker et al., 2011; Soltani et al., 2025; Willette et al., 2015)**. While T2DM makes well documented contributions to cardiovascular disease (including hypertension and atherosclerosis) and such maladies can result in vascular dementia, it is not likely that this results directly from insulin resistance in the conventional sense. In peripheral tissues afflicted with insulin resistance, impairment of insulin signal transduction suppresses the pathways that would normally activate glucose transporter 4 **(Leto and Saltiel, 2012)., 2012; (Khalid et al., 2021; van Gerwen et al., 2023)**. This mechanism does not make a substantial contribution to glucose transport and utilization in the CNS **(Rebelos et al., 2021)**, where glucose metabolism is primarily driven by electrophysiological activity **(Li et al., 2023; Lundgaard et al., 2015)**.

Although insulin resistance may not reduce glucose utilization in the CNS via the same mechanism seen in the periphery, insulin does quantitatively affect electrophysiology, qualifying the peptide as a neuromodulator **(Ferrario and Reagan, 2018; Nunez et al., 2023)**. Such agents, acting less universally and with less necessity than neurotransmitters, nevertheless contribute importantly to synaptic plasticity (long-term potentiation and depression) and homeostasis (synaptic scaling). Insulin generally enhances the release of, and responses to, neurotransmitters in neurons of the cerebral cortex **(Fetterly et al., 2021)**, suggesting that it might stimulate cerebral glucose utilization by stimulating electrophysiological activity.

Insulin-like growth factors (IGFs) signal through a receptor tyrosine kinase that activates downstream pathways shared, at least partially, with insulin, including insulin/IGF-receptor substrates (IRS1 and -2) and cascades activated by phosphatidylinositol-3 kinases (PI3K) and extracellular signal–regulated kinases (Erks) **(Cheng et al., 2000; Thomas and Huganir, 2004; Zheng and Quirion, 2004)**. These pathways converge on targets that regulate synaptic transmission, including modulation of glutamate receptor trafficking and postsynaptic density composition **(Davila et al., 2007; Yue et al., 2022)**. Given the relative abundance of IGFs, especially IGF-1, in the adult CNS, these factors may be even more significant than insulin in situations where insulin/IGF signaling (IIS) is attenuated by pathogenic processes. IGF-1 is synthesized in hippocampal neurons and released in an activity-dependent, synapse-proximal (autocrine) manner, which initiates rapid activation of IGF-1 receptors on the same stimulated dendritic spine to regulate plasticity at the single-spine level **(Tu et al., 2023)**. Moreover, IGF-1 receptors are highly concentrated at synaptic sites, with the majority localized within approximately 50 nm of pre- and postsynaptic structures, thereby facilitating synapse-specific regulation of transmission and plasticity **(Revelo et al., 2014)**.

It has recently been reported that IGF-1 can acutely dampen AMPAR-mediated synaptic currents in neocortical neurons **(Yue et al., 2022)**, thereby raising the interesting possibility that this growth factor would have impacts on excitatory transmission that are distinct from those of insulin. Previous research has demonstrated the effects of IGF-1 on excitatory synaptic transmission in hippocampal neurons **(Ramsey et al., 2005)**. Similarly, **(Gazit et al., 2016)** reported that IGF-1 receptor signaling differentially regulates AMPAR- and NMDAR-mediated transmission, suggesting that IGF-1 may function as a homeostatic modulator that fine-tunes excitatory synaptic strength in neocortical circuits.

We compared the effects of insulin and IGF-1 on glutamatergic responses in neurons derived from the hippocampus and the neocortex. The results indicate that these peptides act in opposing ways with one another, and each acts in an opposing way in the two cell types. These findings are intriguing with regard to the mechanistic details that might explain these phenomena, as well as the conceptual ramifications of two related factors impacting neural circuitry so differently.

## RESULTS

### 1. Insulin enhances glutamate-induced calcium influx in neocortical neurons

To investigate the impact of IGF-1 and insulin on electrophysiological activity, we monitored the impact of these peptides on responses to glutamate receptor activity as reflected by intracellular calcium concentrations ([Ca^2+^]_i_) in neocortical neurons. Neither insulin nor IGF-1 alone exerted a significant effect on [Ca^2+^]_i_. However, a short-term application of insulin resulted in a significantly greater response to glutamate (**Fig. 1**). By contrast, IGF-1 produced a significant decrease in the response.

**Figure 1.**
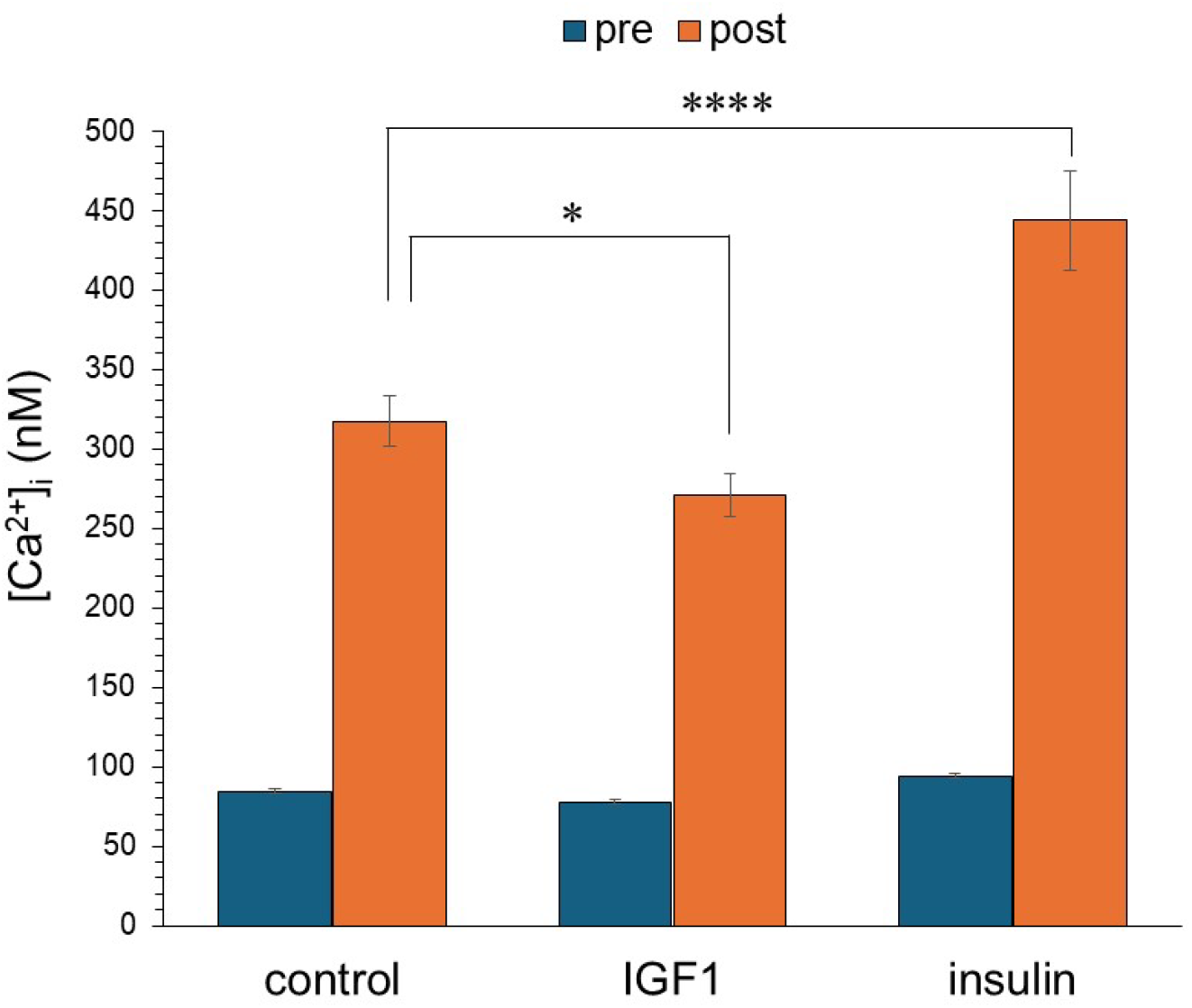
Differential response of insulin and IGF-1 on glutamate-evoked [Ca^2+^]_i_ influx in primary neocortical neurons. Intracellular calcium flux in primary neocortical neurons was measured pre- (without glutamate stimulation) and post- (after glutamate stimulation). In the absence of glutamate, no significant differences in the resting calcium homeostasis were observed among control, IGF-1, or insulin conditions. Following glutamate stimulation, insulin significantly increased calcium influx, compared to control. IGF-1 produced a statistically significant decrease in calcium influx compared to control. Data are presented as mean ± SEM. Statistical comparisons were performed using one-way ANOVA followed by Tukey’s post hoc test. \**P* < 0.05; \*\*\*\**P* < 0.0001 vs. control.

**Figure 2.**
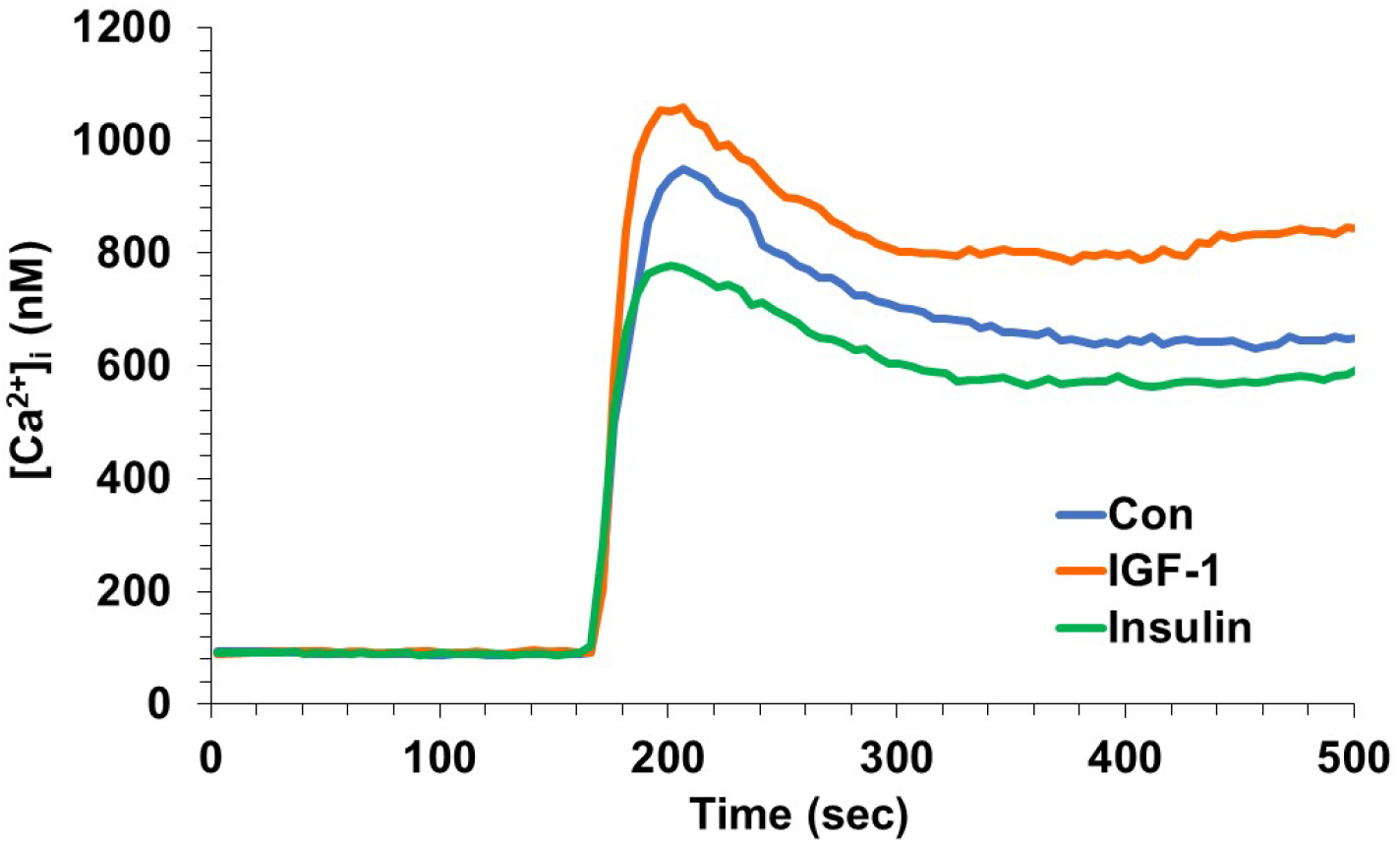
IGF-1 increased glutamate receptor-mediated calcium influx in hippocampal neurons. Hippocampal neurons were treated with IGF-1 (10 ng/mL) or insulin (3 nM), and intracellular calcium influx was measured pre (without glutamate stimulation) and post (after glutamate stimulation). The effects of insulin and IGF-1 on the response to glutamate were significantly different (*P* < 0.0003; 1-way ANOVA and Tukey’s *post hoc*).

### 2. IGF-1 enhances glutamate-induced calcium influx in hippocampal neurons

To determine whether the modulatory effects of IGF-1 and insulin on glutamate receptor-mediated calcium influx are region-specific, analogous experiments were conducted in hippocampal neurons. In contrast to neocortical neurons, hippocampal neurons exhibited a diminution of glutamate responses following treatment with insulin, whereas IGF-1 treatment trended toward an elevated response. Taken together, these findings reveal a spatial and ligand-specific pattern of insulin/IGF signaling.

### 3. Attenuation of glutamate-evoked calcium flux in neocortical neurons by IGF-1 is independent of NMDA receptor activity

One potential explanation for distinct effects of insulin or IGF-1 in different cell types is different balance of contributions from various ionotropic glutamate receptor types to responses in these cell populations. Therefore, we determined the impact of IGF-1 on neocortical neurons’ responses resulting from the NMDAR. CNQX was applied to inhibit AMPA and glycine was also present to increase the opportunity for NMDAR activation. Under these conditions, IGF-1 pretreatment did not affect the initial calcium response, but the sustained elevation was significantly higher in the presence of IGF-1, beginning approximately 45 sec after stimulation **(Fig. 3A)**. Similarly, there was no enhancement by IGF-1 when NMDA itself was used as the agonist (not shown). We next examined the impact of IGF-1 on glutamate-evoked [Ca^2+^]_i_ in neocortical neurons after removal of contributions from NMDARs via application of the NMDAR antagonist 2-amino-5-phosphonopentanoic acid (APV). Under these conditions, IGF-1 still produced an attenuated glutamate response **(Fig. 3B)**. Similar results were obtained after blocking NMDARs with MK-801 (not shown). Together, these findings exclude NMDARs as mediators of the effect of IGF-1 on glutamate responses.

**Figure 3.**
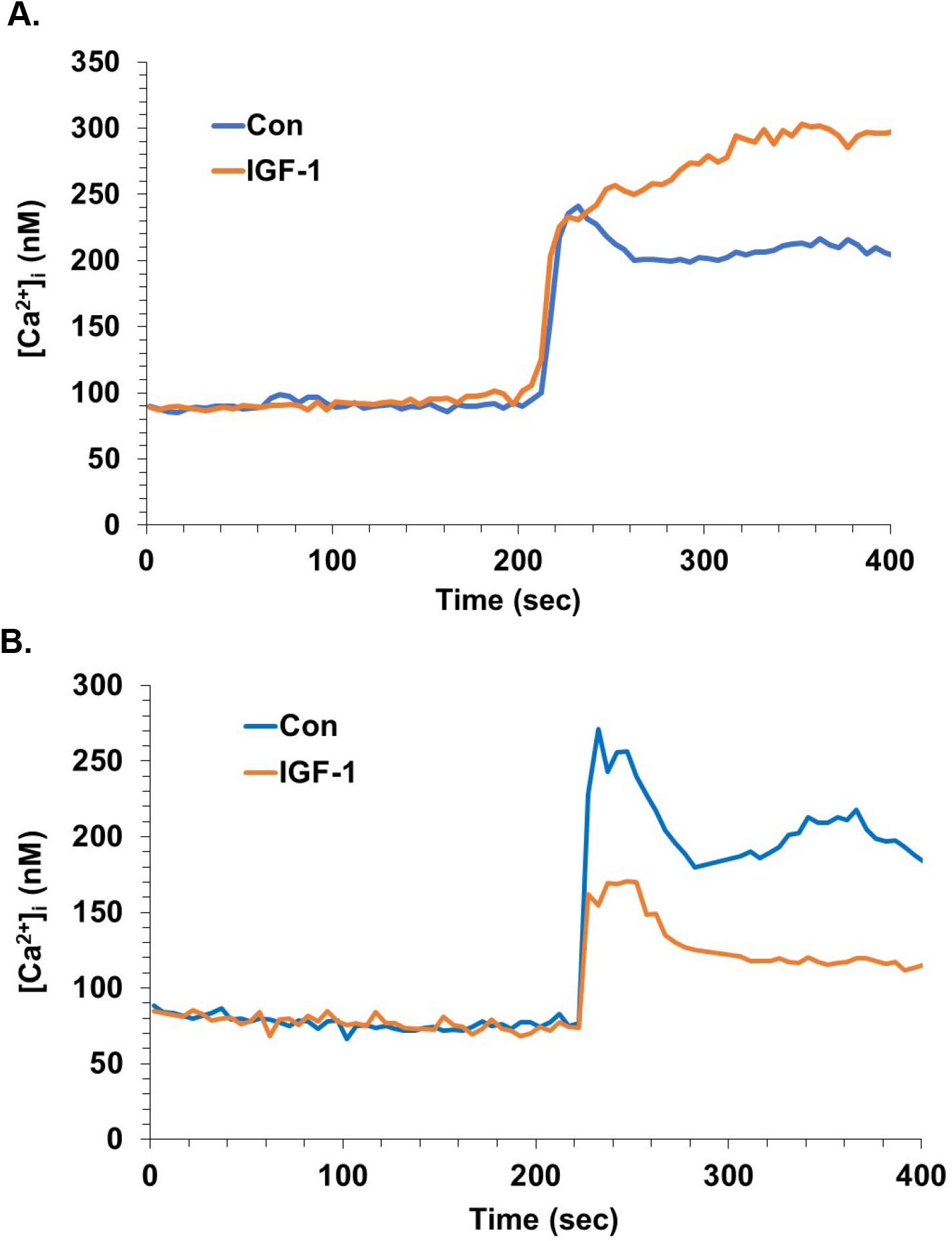
Attenuation of glutamate-evoked calcium flux in neocortical neurons by IGF-1 is independent of NMDAR activity. **A**. Neocortical neurons were pretreated without or with IGF-1 (10 ng/mL), followed by the AMPA receptor antagonist CNQX (125 µM) to isolate the NMDAR response. After 3 minutes, glutamate/glycine (5/10 µM) was applied. The values within a 20-sec window 90 sec after the peak were compared by Student’s *t* test (*P*< 0.012). **B**. Neocortical neurons were monitored for [Ca^2+^]_i_ during exposure to glutamate (5 µM) in the presence of APV (50 µM). One set of cultures was pretreated with IGF-1 (10 ng/mL) for 3 min prior to the glutamate addition. The values within a 20-sec window at the peak were compared by Student’s *t* test (*P*<0.05).

### 4. IGF-1 reduces AMPA receptor-mediated calcium responses in neocortical neurons

Because IGF-1 appeared to elicit an impact on NMDA responses in neocortical neurons that were different from glutamate responses, we explored a role for the other major ionotropic glutamate receptor in neocortical neurons: AMPAR. We tested responses to AMPA itself with and without IGF-1 pretreatment. AMPA application induced a rapid and robust increase in [Ca^2+^]_i_ without IGF-1 (control neurons) (**Fig. 4**), verifying that AMPA receptor–dependent calcium influx is functional even when NMDARs are blocked. When the neocortical neurons were exposed to IGF-1, a significantly attenuated AMPA-evoked calcium response was observed, with a noticeable reduction in both the peak amplitude and sustained phase of the [Ca^2+^]_i_ transient compared with control neurons (*P* < 0.003, Student’s *t* test). This verifies a role for signals initiated at the AMPA receptor in the impact of IGF-1.

**Figure 4.**
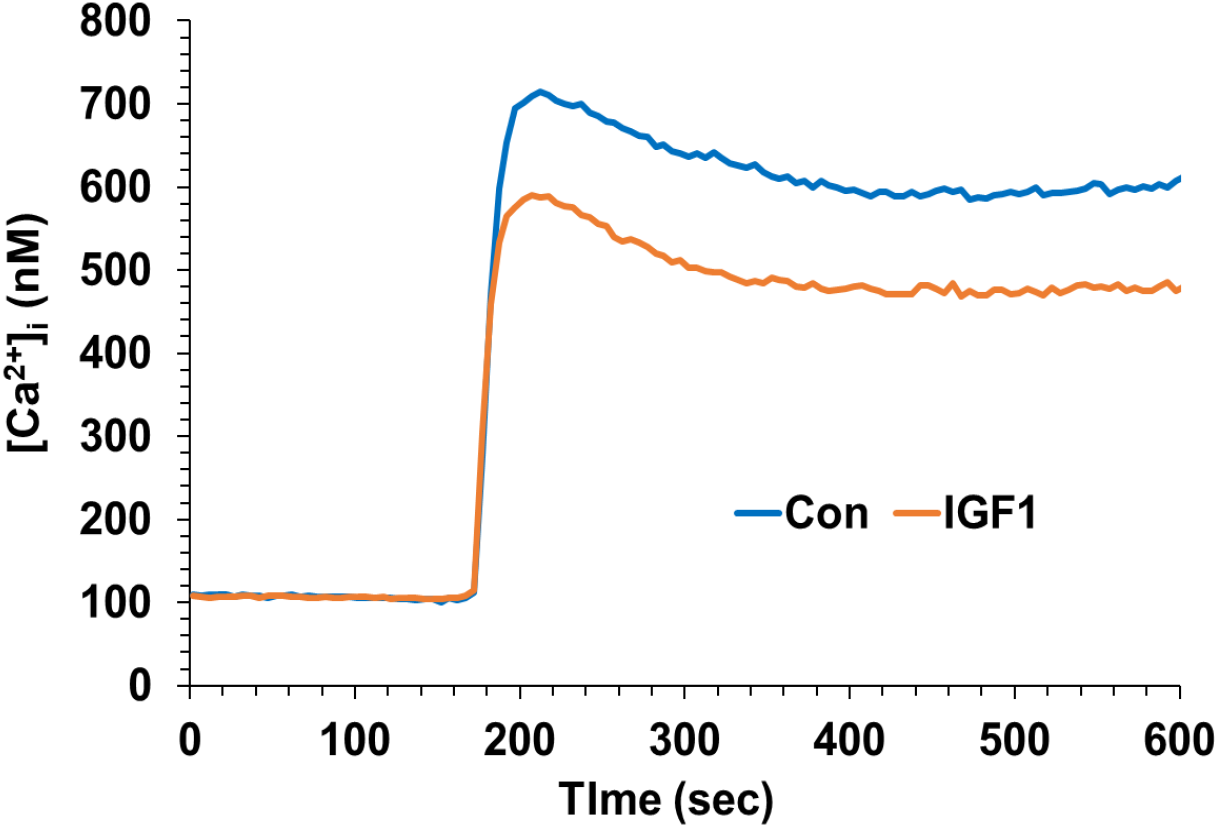
IGF-1 inhibits calcium elevations evoked by AMPA. Neocortical neurons were monitored for [Ca^2+^]_i_ during exposure to AMPA (10 µM) in the presence of APV (50 µM). One set of cultures was pretreated with IGF-1 (10 ng/mL) for 3 min prior to the glutamate addition. The values within a 20-sec window at the peak were compared by Student’s *t* test (*P*<0.003).

### 5. IGF-1 inhibits AMPA receptor-mediated calcium responses independent of L-type calcium channels

While a subset of AMPARs are calcium-permeable, the majority of [Ca^2+^]_i_ elevation by AMPA-receptor stimulation is the result of depolarization and activation of voltage-gated calcium channels (VGCC). It has previously been reported that IGF-1 controls nimodipine-sensitive L-type VGCC in neocortical neurons **(Sanchez et al., 2014)**.

To determine if the inhibition of AMPAR-evoked calcium fluxes by IGF-1 was manifest at the level of L-type VGCC, responses were tested in the presence of nimodipine, an L-type VGCC blocker. Again, NMDARs were blocked with APV (50 µM), and AMPA responses were recorded with or without IGF-1 pretreatment (3 min). Again, the response to AMPA was substantially diminished by IGF-1 (*P* < 0.05, Student’s *t* test) **(Fig. 5)**. These results suggest that IGF-1 modulated calcium responses at some level other than L-type VGCC (potentially, AMPARs themselves).

**Figure 5.**
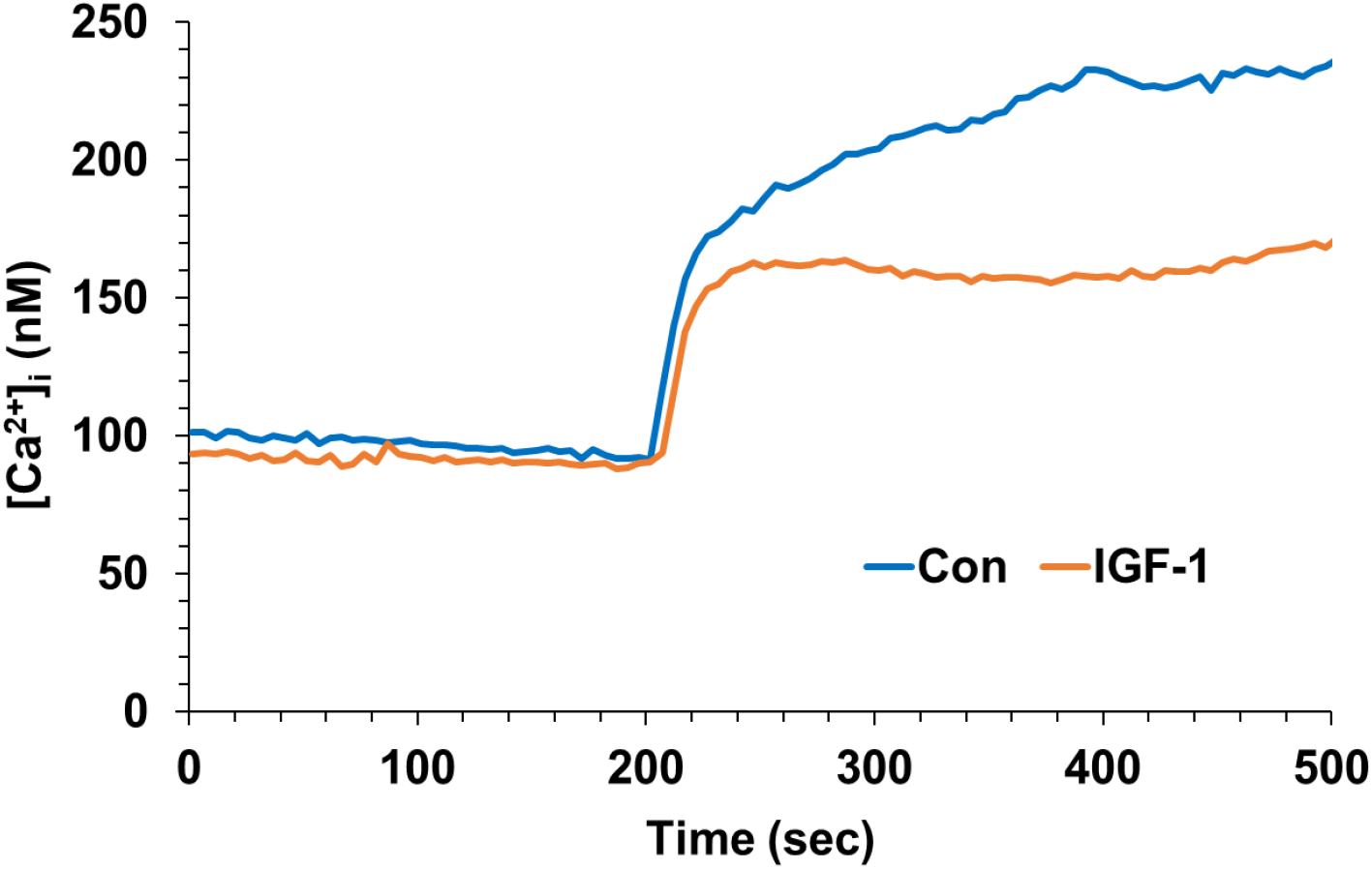
L-type calcium channels are not required for the effect of IGF-1 on AMPA-evoked responses. Cells were pretreated without or with IGF-1 (10 ng/mL) followed by the L-type calcium channel blocker nimodipine (5 µM) and the NMDAR antagonist APV (50 µM). After 3 minutes, they were stimulated with AMPA. Representative traces show that IGF-1 attenuates AMPA-evoked [Ca^2+^]_i_elevation (*P*<0.05; Student’s *t* test) independent of L-type calcium channels.

### 6. Calcium elevations from depolarization alone are unaffected by IGF-1

L-type VGCC were excluded using nimodipine, but it is possible that IGF-1 inhibited the activity of other VGCC triggered by AMPAR-mediated depolarization. To test this possibility, we activated VGCC with KCl to induce membrane depolarization; NMDA and AMPARs were blocked by APV (50 µM) and CNQX (125 µM), respectively. No difference in the KCl-evoked elevation of [Ca^2+^]_i_ was observed between control and IGF-1-treated cells under these conditions (**Fig. 6)**, indicating that IGF-1 did not suppress VGCC generally.

**Figure 6.**
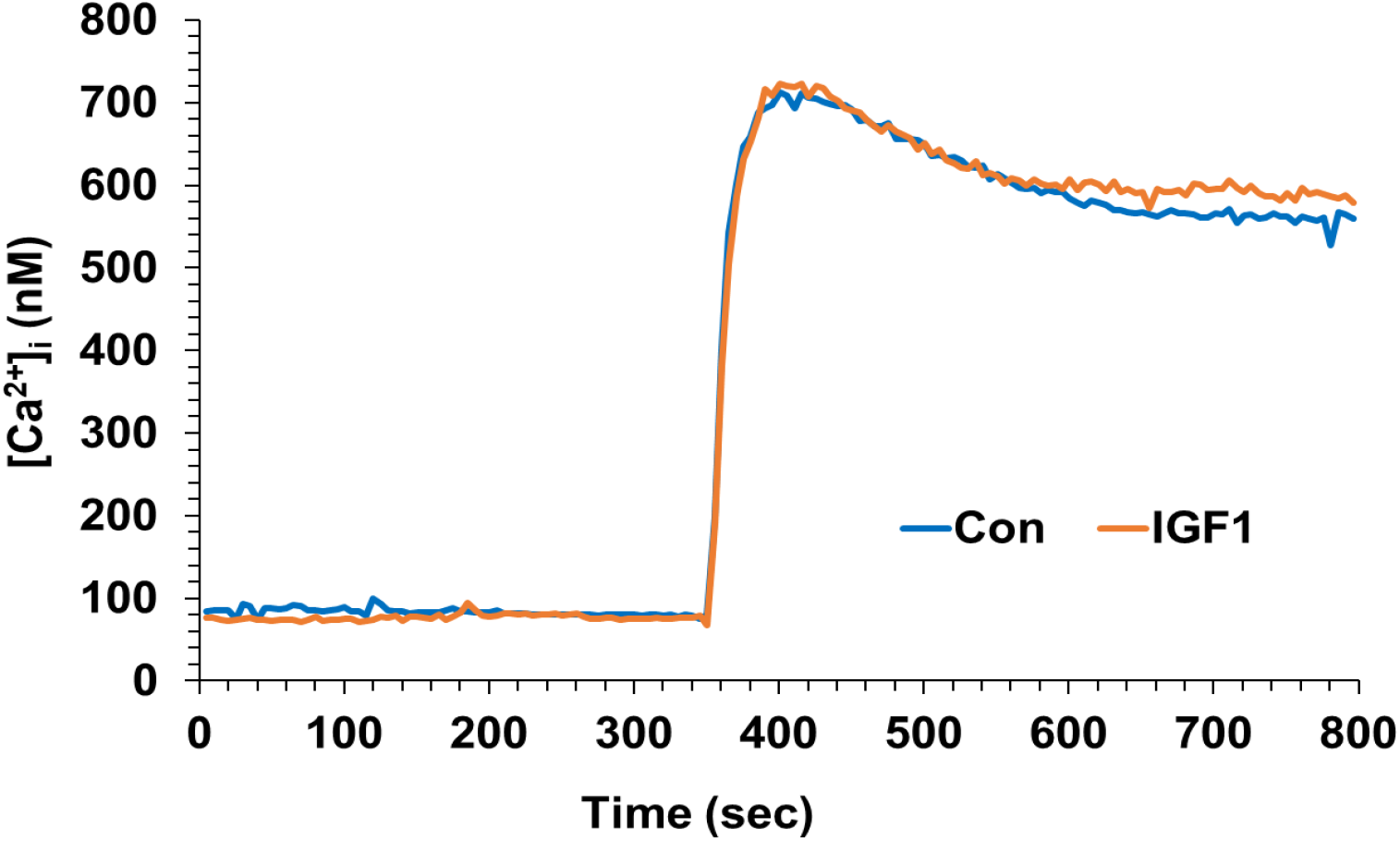
Neurons were pretreated without or with IGF-1 (10 ng/mL), followed by both APV and CNQX (50 µM and 125 µM, respectively) to obviate ionic glutamate receptors. After 3 minutes, the cells were depolarized by raising KCl to 15 mM. The results indicate that IGF-1 does not affect voltage-gated calcium channels directly, suggesting that its direct effect is on AMPARs.

## DISCUSSION

IGF-1 plays a key role in synaptic plasticity, spatial learning, and anxiety-like behavioral processes. These phenomena may be mediated via IGF-1 regulation of neuronal firing and synaptic transmission in many areas of the central nervous system **(Dyer et al., 2016; Garcia-Magro et al., 2025)**. Alongside IGF-1, insulin seems to influence neuronal function and synaptic plasticity through phosphorylation of proteins that control glutamate receptor trafficking, potentiation of NMDAR-mediated responses, and regulation of AMPA receptor endocytosis **(Beattie et al., 2000; Liu et al., 1995; van der Heide et al., 2005)**. Insulin shares downstream signaling pathways with IGF-1, including PI3K/Akt and MAPK cascades, which are critical for neuronal growth and synaptic remodeling **(Fernandez and Torres-Aleman, 2012)**. Disruption of either insulin or IGF-1 signaling in the brain, often referred to as brain insulin resistance and IGF-1 resistance, diminishes these signaling pathways, leading to impaired LTP, altered glutamatergic transmission, and synaptic dysfunction, phenomena increasingly recognized in models of cognitive decline and Alzheimer’s disease **(Ferreira, 2021; Sedzikowska and Szablewski, 2021)**. Given that 1) glucose utilization in the brain is modulated primarily by neurophysiological activity, and 2) insulin and IGF-1 can impact neurophysiological activity, it is possible that these neuromodulators influence cerebral glucose utilization indirectly via their effects on neurotransmission rather than by the glucose transporter 4–dependent mechanisms which operate in peripheral tissues. The effects of IGF-1 on glutamatergic responses in neocortical neurons have not been well elucidated. Various actions of IGF-1 have been reported on calcium or its channels, including some experiments with neocortical neurons **(Sanchez et al., 2014)**. We examined glutamatergic responses to IGF-1 with an emphasis on neocortical neurons. Our results confirm AMPARs as a primary downstream target of IGF-1 signaling and highlight the inhibitory effect of this factor in neurons derived from the neocortex.

By comparing insulin and IGF-1 we confirmed previous reports of opposing effects of these peptides on glutamate receptor activity in hippocampal neurons. Specifically, in hippocampal neurons IGF-1 robustly enhanced glutamate-evoked calcium influx, whereas insulin elicited a modest but consistent suppression of this response. The effect of IGF-1 is consistent with a prior report that acute treatment enhances excitatory synaptic transmission through a postsynaptic mechanism mediated by AMPARs (without involving NMDARs) in hippocampal slices **(Ramsey et al., 2005)**. In addition, insulin has been previously documented to decrease the spontaneous firing rates of hippocampal pyramidal neurons **(Palovcik et al., 1984)**, an effect that likely results from enhanced internalization of GluA2-containing AMPARs **(Beattie et al., 2000; Man et al., 2000)**.

We also tested neurons from the neocortex, which provided confirmatory data for insulin; specifically, elevated responses to glutamate were apparent following insulin. However, effects of IGF-1 on this neuronal type were not well characterized previously, and our findings extended the intriguing opposition between insulin and IGF-1 here as well: insulin elevated and IGF-1 reduced glutamatergic responses. These observations suggest that, despite substantial overlap in their downstream signaling pathways, insulin and IGF-1 exhibit distinct regional specificities that may reflect heterogeneity in distribution and relative expression levels of the primary receptors for these two peptides across neocortical and hippocampal circuits, as well as potential differences in intracellular signaling. Relevant specificities were reported by **(Talbot et al., 2012)**, who found that insulin and IGF-1 had distinct effects on receptors and intracellular signal-transduction elements in normal tissue preparations; moreover, Alzheimer’s disease appeared to create resistance to each of these stimuli differently in hippocampus versus neocortex.

Ionotropic glutamate receptors make the largest contribution to glutamate-evoked calcium fluxes, and NMDARs make the most direct impact via direct calcium conductance through the receptors’ channel. Therefore, we initially performed positive and negative tests of the role of NMDAR in the effects of IGF-1. The stimulation of NMDARs by its namesake agonist was impacted by IGF-1 in a direction opposite that of glutamate stimulation, and glutamate responses were inhibited by IGF-1 when NMDARs were inhibited. This indicates that IGF-1 does not inhibit glutamate responses through suppression of NMDAR, prompting us to examine AMPARs directly. When NMDARs were blocked with APV and AMPA was used as the agonist, IGF-1 produced a decrease in intracellular Ca^2+^ flux, confirming that AMPAR are primarily responsible for the inhibition of glutamate responses by IGF-1. Thus, the dominant effect of IGF-1 on responses to glutamate without blockade of specific receptor subtypes was similar to its effect on AMPAR, indicating that this receptor subtype dominates the response to glutamate in neocortical neurons. Notably, basal Ca^2+^ levels were unaffected by IGF-1, supporting a stimulus-dependent and receptor-specific action of IGF-1.

While a small subset of AMPARs are directly permeable to calcium, most of the calcium flux evoked by AMPAR results from depolarization which opens VGCCs. IGF-1 was previously reported to modulate L-type VGCC **(Blair and Marshall, 1997; Sanchez et al., 2014)**. We tested this possibility in our system via application of the L-type channel blocker nimodipine. The persistence of IGF-1 effects in the presence of nimodipine indicates that L-type VGCC were not involved in the inhibitory effect of IGF-1. However, it is possible that IGF-1 has direct effects on another type of VGCC. Thus, we tested for effects of IGF-1 on calcium fluxes triggered by general membrane depolarization. IGF-1 had no effect on these fluxes. Therefore, this control makes it unlikely that IGF-1 broadly suppresses neuronal excitability or VGCCs directly. Instead, our findings suggest that the effect of IGF-1 involves a direct impact on AMPARs. One hypothesis is that IGF-1 reduces AMPAR surface availability, as supported by the ability of IGF-1 to reduce AMPAR-mediated excitatory synaptic transmission in layer V pyramidal neurons of the adult prefrontal cortex, which aligns with a decrease in membrane localization of GluR1 and -2 **(Yue et al., 2022)**. It is also possible that IGF-1 alters AMPAR conductivity, e.g., via impacts on phosphorylation **(Esteban et al., 2003; Hayashi and Huganir, 2004; Sathler et al., 2021)**. The rapidity of the effect, manifest with as little as 2 minutes of exposure to IGF-1, suggests a mechanism involving post-translational modification such as phosphorylation.

IGF-1 has been shown to modulate synaptic strength and excitatory transmission in hippocampus by regulating AMPAR surface expression and subunit composition, thereby reducing calcium-permeable AMPAR activity without changing basal calcium homeostasis **(Yue et al., 2022)**. These findings, together with the observation that IGF-1 does not suppress NMDAR-mediated calcium responses, indicate that AMPARs represent a primary locus of IGF-1-mediated regulation of excitatory calcium dynamics in neocortical neurons. This suggests that IGF-1 is preferentially integrated into mechanisms governing synaptic strength in neocortical circuits. This may bear some relationship to synaptic scaling, a homeostatic mechanism by which the pro-inflammatory cytokine tumor necrosis factor (TNF) serves to increase AMPAR currents to compensate for circumstances that diminish synaptic activity, typically through an involvement of glial elements in tripartite synaptic units **(Heir et al., 2024; Stellwagen and Malenka, 2006)**. Consequently, IGF-1 may intersect with or compete with TNF-dependent pathways to fine-tune AMPAR trafficking and sustain excitatory synaptic equilibrium. It is also noteworthy that IGF-1 enhances neuronal survival in many circumstances. Given the well-established role of glutamate receptors and calcium in excitotoxicity **(Fairless et al., 2021; Mark et al., 2001)**, it is possible that some portion of IGF-1-mediated neuroprotection involves suppression of glutamate-evoked calcium elevations.

Although the findings provide important insights, there are inherent limitations to this study. The findings are derived from *in vitro* neuronal cultures and may not fully represent *in vivo* physiology. Furthermore, we did not directly assess AMPAR subunit composition, ion conductance, or trafficking mechanisms underlying IGF-1 action.

Our study offers several directions for further research. We have determined that AMPAR are the main targets; further research is needed to identify the precise signaling pathways linking IGF-1 receptor activation to AMPA receptor regulation. Direct mechanistic evidence for how IGF-1 modifies AMPA receptor function could be obtained by measuring AMPA receptor surface expression and phosphorylation states. To characterize their distinct contributions to the observed modulation of calcium flux, pharmacological manipulation of important IGF-1 signaling pathways, such as PI3K/AKT, ERKs, and phospholipase Cγ, would be informative.

## Conclusion

In conclusion, our findings indicate that IGF-1 attenuates excitatory neurotransmission by selectively diminishing the activity of AMPAR in neocortical neurons, independent of NMDAR activation, while maintaining the function of VGCC. Collectively, these findings further describe the actions of IGF-1 as a regulator of excitatory neurotransmission at AMPAR-dependent synapses and suggest that the loss of this modulatory control may contribute to calcium dysregulation and synaptic vulnerability in insulin-resistant and neurodegenerative brain states.

## MATERIALS AND METHODS

### Primary neocortical neuron culture

Primary cultures of neocortical neurons were derived from aseptic dissection of the cerebral cortex of fetal (embryonic day 18) Sprague-Dawley rats using a modification of the previously described method **(Mao et al., 2016)**. In brief, neocortical tissue was stripped of meninges and digested with 1 mg/ml trypsin (Worthington Biochemical) for 4 min, followed by trypsin inhibitor (GIBCO) for 8 min in calcium- and magnesium-free Hank’s balance salt solution (HBSS; GIBCO) buffered to pH 7.2 with HEPES. Cells were mechanically dissociated by trituration with a fire-polished glass Pasteur pipette. The dissociated cells were filtered through a nylon cell strainer (40 µm, Corning) to obtain a single-cell suspension. The cells were seeded onto glass-bottomed 35-mm dishes (MatTek) in minimal essential medium containing 10% FBS, 14.7 mM KCl, and 10 µg/ml gentamycin sulfate. After 2 h to allow attachment, this medium was replaced with Neurobasal-A (NBA) medium (GIBCO) supplemented with 2% B-27 (GIBCO), 500 µM glutamine, and 10 µg/ml gentamycin sulfate. All animal handling was performed in accordance with the guidelines of the National Institutes of Health (NIH) and approved protocols of the Institutional Animal Care and Use Committee (IACUC) at the Central Arkansas Veterans Healthcare System.

### Drug Preparation

In the pharmacological isolation of evoked currents, the drugs used included the AMPA receptor antagonist 6-cyano-7-nitroquinoxaline-2,3-dione (CNQX; Tocris) and nimodipine (Tocris), an L-type calcium channel blocker, dissolved in dimethyl sulfoxide (DMSO) to create a stock solution, ensuring the final DMSO concentration was below 0.05%. Glutamate (Millipore-Sigma), glycine (Fisher), and KCl (Fisher) were all prepared as stock solutions using deionized water. Acidic ligands, specifically NMDAR antagonist 2-amino-5-phosphonovaleric acid (APV) and α-amino-3-hydroxy-5-methyl-4-isoxazolepropionic acid (AMPA; Tocris), were dissolved in water with equinormal OH^-^ counterion. IGF-1 (R&D Systems) was dissolved in PBS with 100 µg/mL fatty acid–free BSA (Roche Diagnostics), divided into aliquots, and stored at −80 °C until needed; insulin (Lilly) was diluted from its liquid stock to a working solution of 600 nM. Throughout the experiments, each drug was administered in a 10 µl volume at the working concentration and were applied using a centripetal mixing method by gentle pipetting (4–5 repetitions) to ensure uniform distributions.

### Calcium measurements

Neurons were employed for ratiometric calcium analyses at days *in vitro* (DIV) 10–14, a developmental stage during which neocortical neurons demonstrate mature synaptic connectivity and stable calcium signaling.

Fura-2 AM (Invitrogen) was prepared in DMSO as a 1 mM stock solution immediately before cell loading. Cells were loaded with 5 μM of (fura-2 AM) for 30 min at 37 °C, followed by a 30-min dye-free chase period to allow de-esterification of the AM group. Immediately before imaging, the existing culture medium was washed once and replaced with a buffer composed of HBSS with 10 mM HEPES and 10 μg/mL gentamycin (pH 7.2).

Single-cell fura-2 AM measurements were made using an Intracellular Imaging Inc. system with a custom-designed filter changer to alternate between 340 and 380 nm excitation. Fluorescent emission intensities of individual cells were measured at 505 nM with an integrated CCD camera, and the emission elicited by 340 nm excitation was divided by that elicited by 380 nm excitation to arrive at a ratio. Calcium levels were calculated by the InCytIm2 software, which interpolated the ratios into a standard curve created by illumination of fura-2 solutions containing known concentrations of Ca^2+^. Cells were imaged under control conditions and in the presence or absence of IGF-1, followed by application of pharmacological blockers and agonists. Basal intracellular Ca^2+^ levels stabilized at ∼55–70 nM and were monitored throughout the recording. Traces represent the average neuronal responses from ≥60 cells in three cultures.

### Statistics

Data sets were analysed by GraphPad Prism using an unpaired two-tailed Student’s *t*-test for comparison between the control and IGF-1-treated group. In experiments with more than two treatments, 1-way ANOVA was performed with Tukey’s *post hoc* test. Statistical significance was defined as *P* < 0.05.

## ACKNOWLEDGMENTS

Supported by grants from the National Institutes of Health (U.S.A.): R01AG071782 and R01AG084473.

